# Genome sequencing reveals Zika virus diversity and spread in the Americas

**DOI:** 10.1101/109348

**Authors:** Metsky H.C., Matranga C.B., Wohl S., Schaffner S.F., Freije C.A., Winnicki S.M., West K., Qu J., Baniecki M.L., Gladden-Young A., Lin A.E., Tomkins-Tinch C.H., Ye S.H., Park D.J., Luo C.Y., Barnes K.G., Shah R.R., Chak B., Barbosa-Lima G., Delatorre E., Vieira Y.R., Paul L.M., Tan A.L., Barcellona C.M., Porcelli M.C., Vasquez C., Cannons A.C., Cone M.R., Hogan K.N., Kopp E.W., Anzinger J.J., Garcia K.F., Parham L.A., Gélvez Ramírez R.M., Miranda Montoya M.C., Rojas D.P., Brown C.M., Hennigan S., Sabina B., Scotland S., Gangavarapu K., Grubaugh N.D., Oliveira G., Robles-Sikisaka R., Rambaut A., Gehrke L., Smole S., Halloran M.E., Villar Centeno L.A., Mattar S., Lorenzana I., Cerbino-Neto J., Valim C., Degrave W., Bozza P.T., Gnirke A., Andersen K.G., Isern S., Michael S.F., Bozza F.A., Souza T.M.L., Bosch I., Yozwiak N.L., MacInnis B.L., Sabeti P.C.

**Author notes:** co-first author (Metsky H.C., Matranga C.B., Wohl S., Schaffner S.F.). co-senior author (Andersen K.G., Isern S., Michael S.F., Bozza F.A., Souza T.M.L., Bosch I., Yozwiak N.L., MacInnis B.L., Sabeti P.C.). co-corresponding author: B.L.M.; T.M.L.S.

## Abstract

Despite great attention given to the recent Zika virus (ZIKV) epidemic in the Americas, much remains unknown about its epidemiology and evolution, in part due to a lack of genomic data. We applied multiple sequencing approaches to generate 110 ZIKV genomes from clinical and mosquito samples from 10 countries and territories, greatly expanding the observed viral genetic diversity from this outbreak. We analyzed the timing and patterns of introductions into distinct geographic regions; our phylogenetic evidence suggests rapid expansion of the outbreak in Brazil and multiple introductions of outbreak strains into Puerto Rico, Honduras, Colombia, other Caribbean islands, and the continental US. We find that ZIKV circulated undetected in multiple regions for many months before the first locally transmitted cases were confirmed, highlighting the importance of viral surveillance. We identify mutations with possible functional implications for ZIKV biology and pathogenesis, as well as those potentially relevant to the effectiveness of diagnostic tests.

---

Since its introduction into the Americas, mosquito-borne ZIKV (Family: *Flaviviridae*) has spread rapidly, causing hundreds of thousands of cases of ZIKV disease, as well as ZIKV congenital syndrome and likely other neurological complications^1-3^. Phylogenetic analysis of ZIKV can reveal the trajectory of the outbreak and detect mutations that may be associated with new disease phenotypes or affect molecular diagnostics. Despite the 70 years since its discovery and the scale of the recent outbreak, however, fewer than 100 ZIKV genomes have been sequenced directly from clinical samples. This is due in part to technical challenges posed by low viral loads (for example, often orders of magnitude lower than in Ebola virus or dengue virus infection^4-6^), and by loss of RNA integrity in samples collected and stored without sequencing in mind. Culturing the virus increases the material available for sequencing but can result in genetic variation that is not representative of the original clinical sample.

We sought to gain a deeper understanding of the viral populations underpinning the ZIKV epidemic by extensive genome sequencing of the virus directly from samples collected as part of ongoing surveillance. We initially pursued unbiased metagenomic RNA sequencing to capture both ZIKV and other viruses known to be co-circulating with ZIKV^5^. In most of the 38 samples examined by this approach there proved to be insufficient ZIKV RNA for genome assembly, but it still proved valuable to verify results from other methods. Metagenomic data also revealed RNA from other viruses, including 41 likely novel viral sequence fragments in mosquito pools (Extended Data Table 1). In one patient we detected no ZIKV sequence but did assemble a complete genome from dengue virus (type 1), one of the viruses that co-circulates with and presents similarly to ZIKV^7^.

To capture sufficient ZIKV content for genome assembly, we turned to two targeted approaches for enrichment before sequencing: multiplex PCR amplification^8^ and hybrid capture^9^. We sequenced and assembled complete or partial genomes from 110 samples from across the epidemic, out of 229 attempted (221 clinical samples from confirmed and possible ZIKV disease cases and eight mosquito pools; Table 1, **Supplementary Table 1**). This dataset, which we used for further analysis, includes 110 genomes produced using multiplex PCR amplification (amplicon sequencing) and a subset of 37 genomes produced using hybrid capture (out of 66 attempted). Because these approaches amplify any contaminant ZIKV content, we relied heavily on negative controls to detect artefactual sequence, and we established stringent, method-specific thresholds on coverage and completeness for calling high confidence ZIKV assemblies (Fig. 1a). Completeness and coverage for these genomes are shown in Fig. 1b and c; the median fraction of the genome with unambiguous base calls was 93%. Per-base discordance between genomes produced by the two methods was 0.017% across the genome, 0.15% at polymorphic positions, and 2.2% for minor allele base calls. Concordance of within-sample variants is shown in more detail in Fig. 1d-f. Patient sample type (urine, serum, or plasma) made no significant difference in sequencing success in our study (Extended Data Fig. 1).

**Table 1.** Samples and genomes by region. Sample source information and sequencing results for 229 clinical and mosquito pool samples. Continental US includes 8 mosquito pool samples; all others are clinical samples. In the final column, genomes generated by both methods are counted only once. “Other” includes regions without a ZIKV genome included in downstream analysis.

| Country or territory | Samples | Samples with metagenomic data | Amplicon sequencing genomes | Hybrid capture genomes | Total genomes |
| --- | --- | --- | --- | --- | --- |
| Brazil | 53 | 12 | 27 | 7 | 27 |
| Colombia | 20 | 0 | 4 | 2 | 4 |
| Dominican Republic | 45 | 7 | 30 | 9 | 30 |
| Guatemala/El Salvador | 3 | 0 | 1 | 0 | 1 |
| Haiti | 4 | 0 | 1 | 0 | 1 |
| Honduras | 20 | 6 | 18 | 8 | 18 |
| Jamaica | 20 | 0 | 5 | 0 | 5 |
| Martinique | 3 | 0 | 1 | 0 | 1 |
| Puerto Rico | 15 | 0 | 3 | 1 | 3 |
| Continental US | 36 | 12 | 20 | 10 | 20 |
| Other | 10 | 1 | 0 | 0 | 0 |
| Total | 229 | 38 | 110 | 37 | 110 |

To investigate the spread of ZIKV in the Americas we performed a phylogenetic analysis of the 110 genomes from our dataset, together with 64 published genomes available on NCBI GenBank and in our companion papers^10,11^ (Fig. 2a). Our reconstructed phylogeny (Fig. 2b), which is based on a molecular clock (Extended Data Fig. 2), is consistent with the outbreak originating in Brazil^12^: Brazil ZIKV genomes appear on all deep branches of the tree, and their most recent common ancestor is the root of the entire tree. We estimate the date of that common ancestor to have been in early 2014 (95% credible interval, CI, August 2013 to July 2014). The shape of the tree near the root remains uncertain (i.e. the nodes have low posterior probabilities) because there are too few mutations to clearly distinguish the branches. This pattern suggests rapid early spread of the outbreak, consistent with the introduction of a new virus to an immunologically naive population. ZIKV genomes from Colombia (*n*=10), Honduras (*n*=18), and Puerto Rico (*n*=3) cluster within distinct, well-supported clades. We also observed a clade consisting entirely of genomes from patients who contracted ZIKV in one of three Caribbean countries (the Dominican Republic, Jamaica, and Haiti) or the continental US, containing 30 of 32 genomes from the Dominican Republic and 19 of 20 from the continental US. We estimated the within-outbreak substitution rate to be 1.15x10^−3^ substitutions/site/year (95% CI [9.78x10^−4^, 1.33x10^−3^]), similar to prior estimates for this outbreak^12^. This is somewhat higher (1.3x–5x) than reported rates for other flaviviruses^13^, but is measured over a short sampling period, and therefore may include a higher proportion of mildly deleterious mutations that have not yet been removed through purifying selection.

**Figure 1.**
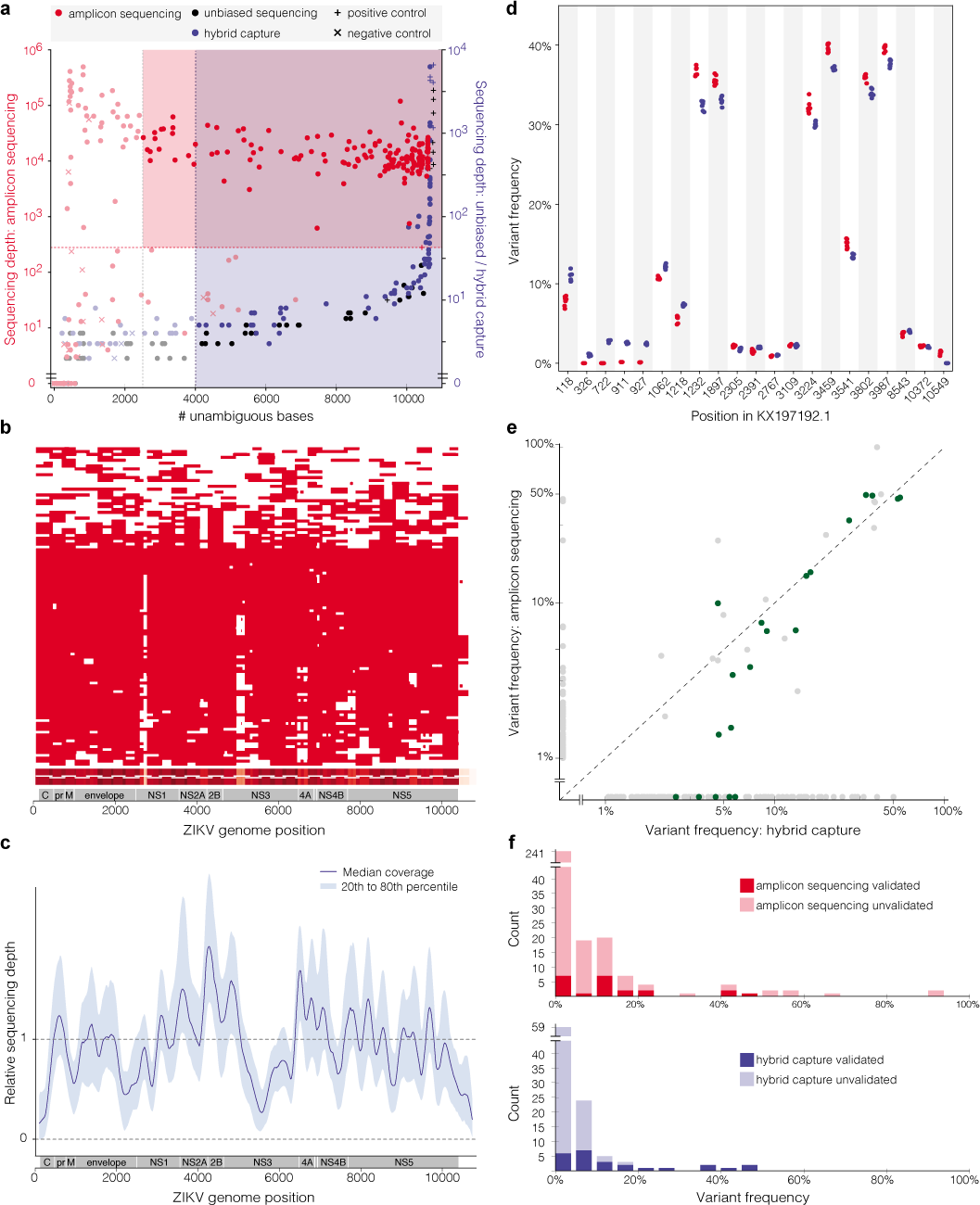
Sequence data from clinical and mosquito samples. **(a)** Thresholds used to select samples for downstream analysis. Each point is a replicate. Red and blue shading: regions of accepted amplicon sequencing and hybrid capture genome assemblies, respectively. Not shown: hybrid capture positive controls with depth >10,000x. **(b)** Amplicon sequencing coverage by sample (row) across the ZIKV genome. Red: sequencing depth ≥100x; heatmap (bottom) sums coverage across all samples. White horizontal lines: amplicon locations. **(c)** Relative sequencing depth across hybrid capture genomes. **(d)** Within-sample variants for a single cultured isolate (PE243) across seven technical replicates. Each point is a variant in a replicate identified using amplicon sequencing (red) or hybrid capture (blue). Variants are plotted if the pooled frequency across replicates by either method is ≥1%. **(e)** Within-sample variant frequencies across methods. Each point is a variant in an individual sample and points are plotted on a log-log scale. Green points: “verified” variants detected by hybrid capture that pass strand bias and frequency filters. **(f)** Counts of within-sample variants across two replicate libraries, for each method. Variants are plotted in the frequency bin corresponding to the higher of c. In (e-f), frequencies <1% are shown at 0%.

Determining when ZIKV arrived in specific regions helps elucidate the spread of the outbreak and track rising incidence of possible complications of ZIKV infection. The majority of the ZIKV genomes from our study fall into four major clades from different geographic regions, for which we estimated a likely date for ZIKV arrival. In each case, the date was months earlier than the first confirmed, locally transmitted case, indicating ongoing local circulation of ZIKV before its detection. In Puerto Rico, the estimated date was 4.5 months earlier than the first confirmed local case^14^; it was 8 months earlier in Honduras^15^, 5.5 months earlier in Colombia^16^, and 9 months earlier for the Caribbean/continental US clade^17^. In each case, the arrival date represents the estimated time to the most recent common ancestor (tMRCA) for the corresponding clade in our phylogeny (Fig. 2c). See Extended Data Fig. 3 and Extended Data Table 2 for details. Similar temporal gaps between the tMRCA of local transmission chains and the earliest detected cases were seen when chikungunya virus emerged in the Americas^18^. We also observed evidence for several introductions of ZIKV into the continental US, and found that sequences from mosquito and human samples collected in Florida cluster together, consistent with the finding of local ZIKV transmission in Florida in a companion paper^11^.

**Figure 2.**
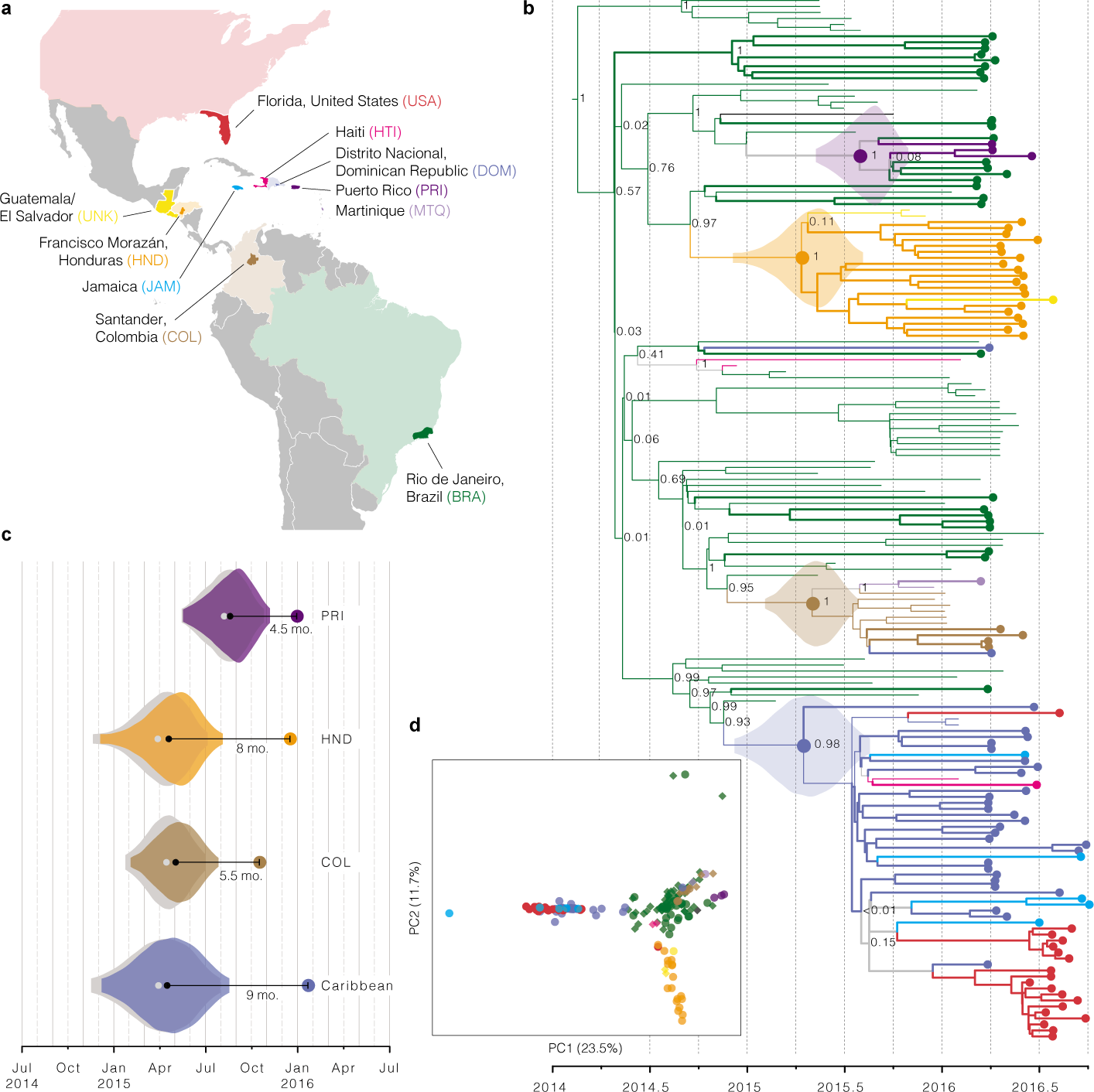
Zika virus spread throughout the Americas. **(a)** Samples were collected in each of the colored countries/ territories. Specific state, department, or province of origin for samples in this study are highlighted if known. **(b)** Maximum clade credibility tree. Dotted tips: genomes generated in this study. Node labels are posterior probabilities indicating support for the node. Violin plots denote probability distributions for the tMRCA of four highlighted clades. **(c)** Time elapsed between estimated tMRCA and date of first confirmed, locally transmitted case. Color: distributions based on relaxed clock model (also shown in (b)); grey: strict clock. “Caribbean” includes the continental US. **(d)** Principal component analysis of variants. Circles: data generated in this study; diamonds: other publicly available genomes from this outbreak. Percentage of variance explained by each component is indicated on axis.

Principal component analysis (PCA) is consistent with the phylogenetic observations (Fig. 2d). It shows tight clustering among ZIKV genomes from the continental US, the Dominican Republic, and Jamaica. ZIKV genomes from Brazil and Colombia are similar and distinct from genomes sampled in other countries. ZIKV genomes from Honduras form a third cluster that also contains genomes from Guatemala or El Salvador. The PCA results show no clear stratification of ZIKV within Brazil.

Genetic variation can provide important clues to understanding ZIKV biology and pathogenesis and can reveal potentially functional changes in the virus. We observed 1030 single nucleotide polymorphisms (SNPs) in the complete dataset, well distributed across the genome (Fig. 3a). Any effect of these mutations cannot be determined from these data; however, the most likely candidates for functional mutations would be among the 202 nonsynonymous SNPs (Supplementary Table 2) and the 32 SNPs in the 5’ and 3’ untranslated regions (UTRs). Adaptive mutations are more likely to be found at high frequency or to be seen multiple times, although both effects can also occur by chance. We observed five positions with nonsynonymous mutations at >5% minor allele frequency that occur on two or more branches of the tree (Fig. 3b); two of these (at 4287 and 8991) occur together and might represent incorrect placement of a Brazil branch in the tree. The remaining three are more likely to represent multiple nonsynonymous mutations; one (at 9240) appears to involve nonsynonymous mutations to two different alleles.

To assess the possible biological significance of these mutations, we looked for evidence of selection in the ZIKV genome. Viral surface glycoproteins are known targets of positive selection, and mutations in these proteins can confer adaptation to new vectors^19^ or aid immune escape^20,21^. We therefore searched for an excess of nonsynonymous mutations in the ZIKV envelope glycoprotein (E). However, the nonsynonymous substitution rate in E proved to be similar to that in the rest of the coding region (Fig. 3c, left); moreover, amino acid changes were significantly more conservative in that region than elsewhere (Fig. 3c, middle and right). Any diversifying selection occurring in the surface protein thus appears to be operating under selective constraint. We also found evidence for purifying selection in the ZIKV 3’ UTR (Fig. 3d, Supplementary Table 3), a region important for viral replication^22^.

While the transition-to-transversion ratio (6.98) was within the range seen in other viruses^23^, we observed a significantly higher frequency of C-to-T and T-to-C substitutions than other transitions (Fig. 3d, Extended Data Fig. 4, Supplementary Table 3). This enrichment is apparent both in the genome as a whole and at 4-fold degenerate sites, where selection pressure is minimal. Many processes may contribute to this conspicuous mutation pattern, including mutational bias of the ZIKV RNA-dependent RNA polymerase, host RNA editing enzymes (e.g. APOBECs, ADARs) acting upon viral RNA, and chemical deamination, but further investigation is required to determine the cause of this phenomenon.

**Figure 3.**
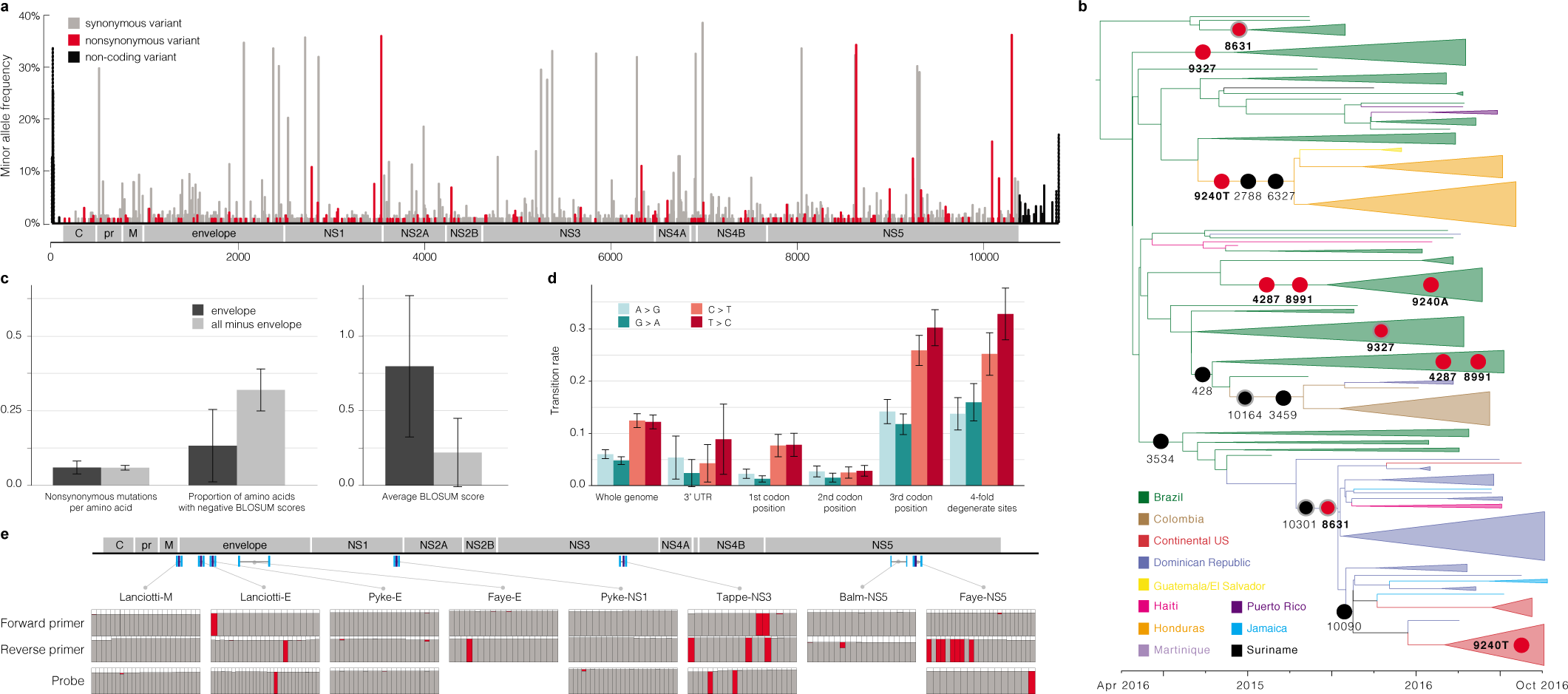
Geographic and genomic distribution of Zika virus variation. **(a)** Location of variants in the ZIKV genome. The minor allele frequency is the proportion of the 174 genomes from this outbreak that share a variant. Dotted bars: <25% of samples had a base call at that position. **(b)** Phylogenetic distribution of nonsynonymous variants with minor allele frequency ≥5%, shown on the branch where the mutation most likely occurred. Grey outline: variant might be on next-most ancestral branch (in two cases, 2 branches upstream), but exact location is unclear because of missing data. Red circles: variants occurring at more than one location in the tree. **(c)** Conservation of the ZIKV envelope (E) region. Left: nonsynonymous variants per amino acid for the E region (dark grey) and the rest of the coding region (light grey). Middle: proportion of nonsynonymous variants resulting in negative BLOSUM62 scores, which indicate unlikely or extreme substitutions (p < 0.039, χ^2^ test). Right: average of BLOSUM62 scores for nonsynonymous variants (p < 0.037, 2-sample *t*-test). **(d)** Constraint in the ZIKV 3’ UTR and transition rates over the ZIKV genome. **(e)** ZIKV diversity in diagnostic primer and probe regions. Top: locations of published probes (dark blue) and primers (cyan)^26^–^31^ on the ZIKV genome. Bottom: each column represents a nucleotide position in the probe or primer. Colors in the column indicate the fraction of ZIKV genomes (out of 174) that match the probe/primer sequence (grey), differ from it (red), or have no data for that position (white).

Mismatches between PCR assays and viral sequence are a potential source of poor diagnostic performance in this outbreak^24^. To assess the potential impact of ongoing viral evolution on diagnostic function, we compared eight published qRT-PCR-based primer/probe sets to our data. We found numerous sites where the probe or primer did not match an allele found among the 174 ZIKV genomes from the current dataset (Fig. 3e). In most cases, the discordant allele was shared by all outbreak samples, presumably because it was present in the Asian lineage that entered the Americas. These mismatches could affect all uses of the diagnostic assay in the outbreak. We also found mismatches from new mutations that occurred following ZIKV entry into the Americas. Most of these were present in less than 10% of samples, although one was seen in 29%. These observations suggest that genome evolution has not caused widespread degradation of diagnostic performance during the course of the outbreak, but that mutations continue to accumulate and ongoing monitoring is needed.

Analysis of within-host viral genetic diversity can reveal important information for understanding virus-host interactions and viral transmission. However, accurately identifying these variants in low-titer clinical samples is challenging, and further complicated by potential artefacts associated with enrichment prior to sequencing. To investigate whether we could reliably detect within-host ZIKV variants in our data, we identified within-host variants in a cultured ZIKV isolate used as a positive control throughout our study, and found that both amplicon sequencing and hybrid capture data produced concordant and replicable variant calls (Fig. 1d). In clinical samples, hybrid capture within-host variants were noisier but contained a reliable subset: although most variants were not validated by the other sequencing method or by a technical replicate, those at high frequency were always replicable, as were those that passed a previously described filter^25^ (Fig. 1e-f, Extended Data Table 3). Within this high confidence set we looked for variants shared between samples as a clue to transmission patterns, but there were too few variants to draw any meaningful conclusions. By contrast, within-host variants identified in amplicon sequencing data were unreliable at all frequencies (Fig. 1f, Extended Data Table 3), suggesting that further technical development is needed before amplicon sequencing can be used to study within-host variation in ZIKV and other clinical samples with low viral titer.

Sequencing low titer viruses like ZIKV directly from clinical samples presents several challenges that have likely contributed to the paucity of genomes available from the current outbreak. While development of technical and analytical methods will surely continue, we note that factors upstream in the process, including collection site and cohort, were strong predictors of sequencing success in our study (Extended Data Fig. 1). This highlights the importance of continuing development and implementation of best practices for sample handling, without disrupting standard clinical workflows, for wider adoption of genome surveillance during outbreaks. Additional sequencing, however challenging, remains critical to ongoing investigation of ZIKV biology and pathogenesis. Together with two companion studies^10,11^, this effort advances both technological and collaborative strategies for genome surveillance in the face of unexpected outbreak challenges.

## Author Contributions

C.B.M., S.W., C.A.F., S.M.W., K.W., J.Q., M.L.B., A.G.-Y., C.Y.L., R.R.S., G.B.-L., Y.R.V., L.M.P., A.L.T., C.M.B., M.C.P., C.Vasquez., A.C.C., M.R.C., K.N.H., E.W.K.IV, J.J.A., K.F.G., L.A.P., R.M.G.R., M.C.M.M., C.M.B., S.H., B.S., S.Scotland., K.G., G.O., R.R.-S., and I.B. performed laboratory work to process samples. H.C.M., C.B.M., C.A.F., S.M.W., K.W., J.Q., M.L.B., C.Y.L., A.G.-Y., N.G.D., A.G., and K.G.A. developed methods for ZIKV detection, targeted enrichment, and/or sequencing library preparation. H.C.M., C.B.M., S.W., S.F.S., M.L.B., A.E.L., C.H.T.-T., S.Y., D.J.P., E.D., A.R., T.M.L.S., I.B., and B.L.M. performed data analyses. S.Smole., L.A.V.C., S.M., I.L., S.I., S.F.M., and F.A.B. led clinical studies and/or study sites. K.G.B., B.C., D.P.R., N.D.G., L.G., M.E.H., A.R., A.G., J.C.-N., C.Valim., W.G., P.T.B., A.G., K.G.A., S.I., S.F.M., F.A.B., T.M.L.S., and I.B. provided critical insights and guidance. H.C.M., C.B.M., T.M.L.S., N.L.Y., B.L.M., and P.C.S. oversaw study design and management. H.C.M., C.B.M., S.W., S.F.S., A.E.L., N.L.Y., B.L.M. and P.C.S. drafted the manuscript. All authors reviewed the manuscript.

### Acknowledgements

We are deeply grateful for the vision and support of Marc and Lynne Benioff, and for the continued support and guidance of Liliana Brown, Eun Mi Lee, and Maria Giovanni (NIAID), and Justine Levin-Allerhand and Eric S. Lander (Broad Institute). We thank Molly Schleicher, Emily Lipscomb, August Felix, Andrea Saltzman, and Stacey Donnelly (Broad Institute) for timely assistance with IRB and Ethics processes; Eva Mair and Liisa Nogelo (Broad Institute), and Erika Carmean (HHMI) for legal counsel; Tamara Mason and the Broad Institute Genomics Platform for sequencing support; Ashley Matthews, Sinéad Chapman, Daniel E. Neafsey, Bruce W. Birren (Broad Institute) for management and guidance; Oliver Pybus (Oxford University) and ZiBRA Project colleagues for sharing data prior to publication; Daniel Olson and Edwin Asturias (Children’s Hospital Colorado) for sharing samples for preliminary studies; Marc Salit (National Institute of Standards and Technology) and Etienne Simon-Loriere (Pasteur Institute) for sharing reagents; and Edward Holmes (University of Sydney), Gonzalo Bello (Fiocruz), Ryan Tewhey, Anne Piantadosi, Chris Edwards and the Sabeti Lab (Broad Institute) for many helpful discussions and critical reading of the manuscript. We are indebted to Zika patients and clinical teams for making this work possible.

## Methods

### Ethics statement

The clinical studies from which samples were obtained were evaluated and approved by relevant Institutional Review Boards/Ethics Review Committees at: Hospital General de la Plaza de la Salud (Santo Domingo, Dominican Republic), University of the West Indies (Kingston, Jamaica), Universidad Nacional Autónoma de Honduras (Tegucigalpa, Honduras), Oswaldo Cruz Foundation (Rio de Janeiro, Brazil), Centro de Investigaciones Epidemiologicas - Universidad Industrial de Santander (Bucaramanga, Colombia), Massachusetts Department of Public Health (Jamaica Plain, Massachusetts), and Florida Department of Health (Tallahassee, Florida). Informed consent was obtained from all participants enrolled in studies at Hospital General de la Plaza de la Salud, Universidad Nacional Autónoma de Honduras, Oswaldo Cruz Foundation, and Universidad Industrial de Santander. IRBs at the University of West Indies, Massachusetts Department of Public Health, and Florida Department of Health granted waivers of consent given this research with leftover clinical diagnostic samples involved no more than minimal risk. Harvard University and Massachusetts Institute of Technology (MIT) Institutional Review Boards/Ethics Review Committees provided approval for sequencing and secondary analysis of samples collected by the aforementioned institutions.

### Sample collections and study subjects

Suspected ZIKV cases (including high-risk travelers) were enrolled through study protocols at multiple aforementioned collection sites. Clinical samples (including blood, urine, cerebrospinal fluid, and saliva) were obtained from suspected or confirmed ZIKV cases and from high-risk travelers. De-identified information about study participants and other sample metadata are reported in Supplementary Table 1.

### Viral RNA isolation

RNA was isolated following manufacturer’s standard operating protocol for 0.14 mL up to 1 mL samples^32^ using the QIAamp Viral RNA Minikit (Qiagen), except that in some cases 0.1 M final concentration of β-mercaptoethanol (as a reducing agent) or 40 μg/mL final concentration of linear acrylamide (Ambion) (as a carrier) were added to AVL buffer prior to inactivation. Extracted RNA was resuspended in AVE buffer or nuclease-free water. In some cases, viral samples were concentrated using Vivaspin-500 centrifugal concentrators (Sigma-Aldrich) prior to inactivation and extraction. In these cases, 0.84 mL of sample was concentrated to 0.14 mL by passing through a 30 kDa filter and discarding the flow through.

### Carrier RNA and host rRNA depletion

In a subset of human samples, carrier poly(rA) RNA and host rRNA were depleted from RNA samples using RNase H selective depletion^9,33^. Briefly, oligo d(T) (40 nt long) and/or DNA probes complementary to human rRNA were hybridized to the sample RNA. The sample was then treated with 15 units of Hybridase Thermostable RNase H (Epicentre) for 30 minutes at 45°C. The complementary DNA probes were removed by treating each reaction with an RNase-free DNase (Qiagen) according to the manufacturer’s protocol. Following depletion, samples were purified using 1.8x volume AMPure RNAclean beads (Beckman Coulter Genomics) and eluted into 10 μl water for cDNA synthesis.

### Illumina library construction and sequencing

cDNA synthesis was performed as described in previously published RNA-seq methods^9^. To track potential cross-contamination, 50 fg of synthetic RNA (gift from M. Salit, NIST) was spiked into samples using unique RNA for each individual ZIKV sample. ZIKV negative control cDNA libraries were prepared from water, human K-562 total RNA (Ambion), or EBOV (KY425633.1) seed stock; ZIKV positive controls were prepared from ZIKV Senegal (isolate HD78788) or ZIKV Pernambuco (isolate PE243; KX197192.1) seed stock. The dual index Accel-NGS® 2S Plus DNA Library Kit (Swift Biosciences) was used for library preparation. Approximately half of the cDNA product was used for library construction, and indexed libraries were generated using 18 cycles of PCR. Each individual sample was indexed with a unique barcode. Libraries were pooled at equal molarity and sequenced on the Illumina HiSeq 2500 or MiSeq (paired-end reads) platforms.

### Amplicon-based cDNA synthesis and library construction

ZIKV amplicons were prepared as described^8,11^, similarly to “RNA jackhammering” for preparing low input viral samples for sequencing^34^, with slight modifications. After PCR amplification, each amplicon pool was quantified on a 2200 Tapestation (Agilent Technologies) using High Sensitivity D1000 ScreenTape (Agilent Technologies). 2 μL of a 1:10 dilution of the amplicon cDNA was loaded and the concentration of the 350-550 bp fragments was calculated. The cDNA concentration, as reported by the Tapestation, was highly predictive of sequencing outcome (i.e. whether a sample passes genome assembly thresholds) (Extended Data Fig. 5). cDNA from each of the two amplicon pools were mixed equally (10-25 ng each) and libraries were prepared using the dual index Accel-NGS® 2S Plus DNA Library Kit (Swift Biosciences) according to manufacturer’s protocol. Libraries were indexed with a unique barcode using 7 cycles of PCR, pooled equally and sequenced on the Illumina MiSeq (250 bp paired-end reads) platform. Primer sequences were removed by hard trimming the first 30 bases for each insert read prior to analysis.

### Zika virus hybrid capture

Viral hybrid capture was performed as previously described^9^. Probes were created to target ZIKV and chikungunya virus (CHIKV). Candidate probes were created by tiling across publicly available sequences for ZIKV and CHIKV on NCBI GenBank^35^. Probes were selected from among these candidate probes to minimize the number used while maintaining coverage of the observed diversity of the viruses. Alternating universal adapters were added to allow two separate PCR amplifications, each consisting of non-overlapping probes. (To download probe sequences, see Supplementary Information.)

The probes were synthesized on a 12k array (CustomArray). The synthesized oligos were amplified by two separate emulsion PCR reactions with primers containing T7 RNA polymerase promoter. Biotinylated baits were in vitro transcribed (MEGAshortscript, Ambion) and added to prepared ZIKV libraries. The baits and libraries were hybridized overnight (∼16 hrs), captured on streptavidin beads, washed, and re-amplified by PCR using the Illumina adapter sequences. Capture libraries were then pooled and sequenced. In some cases, a second round of hybrid capture was performed on PCR-amplified capture libraries to further enrich the ZIKV content of sequencing libraries (Extended Data Fig. 6). In the main text, “hybrid capture” refers to a combination of hybrid capture sequencing data and data from the same libraries without capture (unbiased), unless explicitly distinguished.

### Genome assembly

We assembled reads from all sequencing methods into genomes using viral-ngs v1.13.3^36,37^. We taxonomically filtered reads from amplicon sequencing against a ZIKV reference, KU321639.1. We filtered reads from other approaches against the list of accessions provided in Supplementary Information. To compute results on individual replicates, we *de novo* assembled these and scaffolded against KU321639.1. To obtain final genomes for analysis, we pooled data from multiple replicates of a sample, *de novo* assembled, and scaffolded against KX197192.1. For all assemblies, we set the viral-ngs ‘assembly_min_length_fraction_of_reference’ and ‘assembly_min_unambig’ parameters to 0.01. For amplicon sequencing data, unambiguous base calls required at least 90% of reads to agree in order to call that allele (‘major_cutoff’ = 0.9); for hybrid capture data, we used the default threshold of 50%. We modified viral-ngs so that calls to GATK’s UnifiedGenotyper set ‘min_indel_count_for_genotyping’ to 2.

At 3 sites with insertions or deletions (indels) in the consensus genome CDS, we corrected the genome using Sanger sequencing of the RT-PCR product (namely, at 3447 in the genome for sample DOM_2016_BB-0085-SER; at 5469 in BRA_2016_FC-DQ12D1-PLA; and at 6516-6564 in BRA_2016_FC-DQ107D1-URI, with coordinates in KX197192.1). At other indels in the consensus genome CDS, we replaced the indel with ambiguity.

Depth of coverage values from amplicon sequencing include read duplicates. In all other cases, we removed duplicates with viral-ngs.

### Identification of non-ZIKV viruses in samples by unbiased sequencing

Using Kraken v0.10.6^38^ in viral-ngs, we built a database that includes its default “full” database (which incorporates all bacterial and viral whole genomes from RefSeq^39^ as of October 2015). Additionally, we included the whole human genome (hg38), genomes from PlasmoDB^40^, sequences covering mosquito genomes (*Aedes aegypti*, *Aedes albopictus*, *Anopheles albimanus*, *Anopheles quadrimaculatus*, *Culex quinquefasciatus*, and the outgroup *Drosophila melanogaster*) from GenBank^35^, protozoa and fungi whole genomes from RefSeq, SILVA LTP 16s rRNA sequences^41^, and all sequences from NCBI’s viral accession list^42^ (as of October 2015) for viral taxa that have human as a host. (To download database, see Supplementary Information.)

For each sample, we ran Kraken on data from unbiased sequencing replicates (not including hybrid capture data) and searched its output reports for viral taxa with more than 100 reported reads. We manually filtered the results, removing ZIKV, bacteriophages, and known lab contaminants. For each sample and its associated taxa, we assembled genomes using viral-ngs as described above; results are in Extended Data Table 1a. We used the following genomes for taxonomically filtering reads and as the reference for assembly: KJ741267.1 (cell fusing agent virus), AY292384.1 (deformed wing virus), NC_001477.1 (dengue virus type 1), LC164349.1 (JC polyomavirus). When reporting sequence identity of an assembly to its taxon, we used BLASTN^43^ to determine the identity between the sequence and the reference used for its assembly.

To focus on metagenomics of mosquito pools (Extended Data Table 1b), we considered unbiased sequencing data from 8 mosquito pools (not including hybrid capture data). We first ran the depletion pipeline of viral-ngs on raw data and then ran the viral-ngs Trinity^44^ assembly pipeline on the depleted reads to assemble them into contigs. We pooled contigs from all mosquito pool samples and identified all duplicate contigs with sequence identity >95% using CD-HIT45. Additionally, we used predicted coding sequences from Prodigal 2.6.3^46^ to identify duplicate protein sequences at >95% identity. We classified contigs using BLASTN^43^ against nt and BLASTX^43^ against nr (as of February 2017) and discarded all contigs with an e-value greater than 1E-4. We define viral contigs as contigs that hit a viral sequence, and we manually removed all reverse-transcriptase-like contigs due to their similarity to retrotransposon elements within the *Aedes aegypti* genome. We categorized viral contigs with less than 80% amino acid identity to their best hit as likely novel viral contigs. **Supplementary Table 4** lists the unique viral contigs we found, their best hit, and information scoring the hit.

### Relationship between metadata and sequencing outcome

To determine if available sample metadata are predictive of sequencing outcome, we tested the following variables: sample collection site, patient gender, patient age, sample type, and the number of days between symptom onset and sample collection (“collection interval”). To describe sequencing outcome of a sample *S*, we used the following response variable *Y*_*S*_:

> mean({ I(*R*) * (number of unambiguous bases in *R*) for all amplicon sequencing replicates *R* of *S* }),
>
> where I(*R*)=1 if median depth of coverage of *R* ≥275 and I(*R*)=0 otherwise

This value is listed in **Supplementary Table 1** under “Dependent variable used in regression on metadata”. We excluded the saliva, cerebrospinal fluid, and whole blood sample types due to sample number (*n*=1), and also excluded mosquito pool samples and rows with missing values. We excluded samples from one collection site (prefix “JAM_2016_WI-”) because most had missing values. We treated samples with type “Plasma EDTA” as having type “Plasma”. We treated the “collection interval” variable as categorical (0-1, 2-3, 4-6, and 7+ days).

With a single model we underfit the zero counts, possibly because many zeros (samples without a replicate that passes ZIKV assembly) are truly ZIKV-negative. We thus view the data as coming from two processes: one determining whether a sample is ZIKV-positive or ZIKV-negative, and another that determines, among the observed passing samples, how much of a ZIKV genome we are able to sequence. We modeled the first process, predicting whether a sample is passing, with logistic regression (in R using GLM^47^ with binomial family and logit link); here, the observed passing samples are the samples *S* for which *Y*_*S*_ ≥ 2500. For the second, we performed a beta regression, using only the observed passing samples, of *Y*_*S*_ divided by ZIKV genome length on the predictor variables. We implemented this in R using the betareg package^48^ and transformed fractions from the closed unit interval to the open unit interval as the authors suggest.

To test the significance of predictor variables, we used a likelihood ratio test. For variable *X*_*i*_ we compared a full model (with all predictors) against a model that uses all predictors except *X*_*i*_. Results of these tests are shown in Extended Data Fig. 1a and d. We explore the effects of sample type and collection interval on obtaining a passing assembly in Extended Data Fig. 1b and c, respectively. Error bars are 95% confidence intervals derived from binomial distributions. We explore the effects of these same two variables on *Y*_*S*_ (in passing samples only) in Extended Data Fig. 1e **and** f.

### Criteria for pooling across replicates

We attempted to sequence one or more replicates of each sample and attempted to assemble a genome from each replicate. We discarded data from any replicates whose assembly showed high sequence similarity, in any part of the genome, to our assembly of the genome in a sample consisting of an African (Senegal) lineage (strain HD78788) of ZIKV. We used this sample as a positive control throughout this study, and considered its presence in the assembly of a clinical or mosquito pool sample to be evidence of contamination. Similarly, we discarded data from four replicates belonging to samples from the Dominican Republic because they yielded assemblies that were unexpectedly identical or highly similar to our assembly of the ZIKV isolate PE243 genome, another positive control used in this study. We also discarded data from replicates that showed evidence of contamination, at the RNA stage, by the baits used in hybrid capture; we detected these by looking for adapters that were added to these probes for amplification.

For amplicon sequencing, we consider an assembly of a replicate to be “passing” if it contains at least 2500 unambiguous base calls and has a median depth of coverage of at least 275x over its unambiguous bases (depth includes duplicate reads). For the unbiased and hybrid capture approaches, we consider an assembly of a replicate “passing” if it contains at least 4000 unambiguous base calls. For each approach, the unambiguous base threshold is based on an observed density of negative controls below the threshold (Fig. 1a). For amplicon sequencing assemblies, we added a coverage depth threshold because coverage depth was roughly binary across replicates, with negative controls falling in the lower class. Based on these thresholds, 0 of 99 negative controls used throughout our sequencing runs yield passing assemblies and 32 of 32 positive controls yield passing assemblies.

We consider a sample to have a passing assembly if any of its replicates, by either method, yields an assembly that passes the above thresholds. For each sample with at least one passing assembly, we pooled read data across replicates for each sample, including replicates with assemblies that do not pass the assembly thresholds. When data was available from both amplicon sequencing and unbiased/hybrid capture approaches, we pooled amplicon sequencing data separately from data produced by the unbiased and hybrid capture approaches, the latter two of which were pooled together (henceforth, the “hybrid capture” pool). We then assembled a genome from each set of pooled data. When assemblies on pooled data were available from both approaches, we selected for downstream analysis the assembly from the hybrid capture approach if it had more than 10267 unambiguous base calls (95% of the reference genome used, GenBank accession KX197192.1); when this condition was not met, we selected the one that had more unambiguous base calls.

The number of ZIKV genomes publicly available prior to this study is the result of an NCBI GenBank^35^ search for ZIKV in February 2017. We filtered any sequences with length <4000 nt, excluded sequences that are being published as part of this study or a companion paper^10,11^, excluded sequences from non-human hosts, and excluded sequences labeled as having been passaged. We counted fewer than 100 sequences, the precise number depending on details of the count.

### Visualization of coverage depth across genomes

For amplicon sequencing data, we plotted coverage across the 110 samples that yielded a passing assembly by amplicon sequencing (Fig. 1b). With viral-ngs, we aligned depleted reads to the reference sequence KX197192.1 using the novoalign aligner with options ‘-r Random -1 40 -g 40 -x 20 -t 100 -k’. Because of the nature of amplicon sequencing, duplicates were not identified or removed. We binarized depth at each nucleotide position, showing red if depth of coverage is at least 100x. Rows (samples) are hierarchically clustered to ease visualization.

For hybrid capture sequencing data, we plotted depth of coverage across the 37 samples that yielded a passing assembly (Fig. 1c). We aligned reads as described above for amplicon sequencing data, except we removed duplicates. For each sample, we calculated depth of coverage at each nucleotide position. We then scaled the values for each sample so that each would have a mean depth of 1.0. At each nucleotide position, we calculated the median depth across the samples, as well as the 20^th^ and 80^th^ percentiles. We plotted the mean of each of these metrics within a 200 nt sliding window.

### Multiple sequence alignments

We aligned ZIKV consensus genomes using MAFFT v7.221^49^ with the following parameters: ‘--maxiterate 1000 --ep 0.123 --localpair’.

In Supplementary Data, we provide sequences and alignments used in analyses.

### Analysis of within- and between-sample variants

To measure overall per-base discordance between consensus genomes produced by amplicon sequencing and hybrid capture, we considered all sites where base calls were made in both the amplicon sequencing and hybrid capture consensus genomes of a sample, and we calculated the fraction in which the bases were not in agreement. To measure discordance at polymorphic sites, we took all of the consensus genomes generated in this study that we selected for downstream analysis and searched for positions with polymorphism (see **Criteria for pooling across replicates** for choosing among the amplicon sequencing and hybrid capture genome when both are available). We then looked at these positions in genomes that were available from both methods, and we calculated the fraction in which the alleles were not in agreement.

To measure discordance at minor alleles, we took all of the consensus genomes generated in this study that we selected for downstream analysis and searched for minor alleles. We then looked at all sites at which there was a minor allele and for which genomes from both methods were available, and we calculated the fraction in which the alleles were not in agreement. For these calculations, we tolerated partial ambiguity (e.g. ‘Y’ is concordant with ‘T’). If one genome had full ambiguity (‘N’) at a position and the other genome had an indel, we counted the site as discordant; otherwise, if one genome had full ambiguity, we did not count the site.

After assembling genomes, we determined within-sample allele frequencies for each sample by running V-Phaser 2.0 via viral-ngs^37^ on all pooled reads mapping to the sample assembly. When determining per-library allele counts at each variant position, we modified viral-ngs to require a minimum base (Phred) quality score of 30 for all bases, discard anomalous read pairs, and use per-base alignment quality (BAQ) in its calls to SAMtools^50^ mpileup. This is particularly helpful for filtering spurious amplicon sequencing variants because all generated reads start and end at a limited number of positions (due to the pre-determined tiling of amplicons across the genome). Because amplicon sequencing libraries were sequenced using 250 bp paired-end reads, bases near the middle of the ∼450 nt amplicons fall at the end of both paired reads, where quality scores drop and incorrect base calls are more likely. To determine the overall frequency of each variant in a sample, we summed allele counts (calculated using SAMtools^50^ mpileup via viral-ngs) across libraries.

When comparing variant frequencies between amplicon sequencing (7 technical replicates) and hybrid capture (7 technical replicates) replicates of the PE243 positive control (Fig. 1d), we include only positions at which the mean (pooled) frequency across replicates within at least one method was ≥1%. When comparing allele frequencies between replicate libraries, we restricted the sample set to only samples with a passing assembly in both methods, and included only samples with two or more replicates. In contrast, when comparing alleles across methods we included samples that have a passing assembly by either method, with any number of replicates. For these comparisons, we only included positions with a minor variant; i.e. positions for which both libraries/methods had an allele at 100% were removed, even if the single allele differed between the two libraries/methods. Additionally, we considered any allele with frequency <1% as not found (0%).

When comparing allele frequencies across methods: let *f*_*a*_ and *f*_*hc*_ be frequencies in amplicon sequencing and hybrid capture, respectively. If both are non-zero, we only included an allele if the read depth at its position was ≥1/min(*f*_*a*_, *f*_*hc*_) in both methods, and if depth at the position was at least 100 for hybrid capture and 275 for amplicon sequencing. If *f*_*a*_=0, we required a read depth of max(1/*f*_*hc*_, 275) at the position in the amplicon sequencing method; similarly, if *f*_*hc*_=0 we required a read depth of max(1/*f*_*a*_, 100) at the position in the hybrid capture method. This was to eliminate lack of coverage as a reason for discrepancy between two methods. When comparing allele frequencies across sequencing replicates within a method, we imposed only a minimum read depth (275x for amplicon sequencing and 100x for hybrid capture), but required this depth in both libraries. In samples with more than two replicates, we only considered the two replicates with the highest depth at each plotted position.

We considered allele frequencies from hybrid capture sequencing “verified” if they passed the strand bias and frequency filters described in Gire et al. 2014^25^, with the exception that we imposed a minimum allele frequency of 1% and allowed a variant identified in only one library if its frequency was ≥5%. In Extended Data Table 3 and Fig. 1f, we considered variants “validated” if they were present at ≥1% frequency in both libraries or methods. When comparing two libraries for a given method *M* (amplicon sequencing or hybrid capture): the proportion unvalidated is the fraction, among all variants in *M* at ≥1% frequency in at least one library, of the variants that are at ≥1% frequency in exactly one of the two libraries. Similarly, when comparing methods: the proportion unvalidated for a method *M* is the fraction, among all variants at ≥1% frequency in *M*, of the variants that are at ≥1% frequency in *M* and <1% frequency in the other method.

We initially called SNPs on the aligned consensus genomes using Geneious version 9.1.7^51^. We converted all fully or partially ambiguous calls, which are treated by Geneious as variants, into missing data. We then removed all sites that were no longer polymorphic from the SNP set and re-calculated allele frequencies. A nonsynonymous SNP is shown on the tree (Fig. 3b) if it includes an allele that is nonsynonymous relative to the ancestral state (see **Molecular clock phylogenetics and ancestral state reconstruction** section below) and has a minor allele frequency of >5%; all occurrences of nonsynonymous alleles are shown. (Two SNPs, at positions 2853 and 7229, had nominal derived allele frequencies over 95%; in both cases, the “ancestral” allele was seen only in a small clade within the tree, suggesting that the ancestral allele was incorrectly assigned.) We placed mutations at a node such that the node leads only to samples with the mutation or with no call at that site.

Uncertainty in placement occurs when a sample lacks a base call for the corresponding SNP; in this case, we placed the SNP on the most recent branch for which we have available data. We also used this ancestral ZIKV state to count the frequency of each type of substitution over various regions of the ZIKV genome, per number of available bases in each region (Fig. 3d and Supplementary Table 3).

We quantified the effect of nonsynonymous SNPs using the original BLOSUM62 scoring matrix for amino acids^52^, in which positive scores indicate conservative amino acid changes and negative scores unlikely or extreme substitutions. We assessed statistical significance for equality of proportions by χ2 test (Fig. 3c, middle), and for difference of means by 2-sample *t*-test with Welch-Satterthwaite approximation of df (Fig. 3c, right). Error bars are 95% confidence intervals derived from binomial distributions (Fig. 3c, left and middle; Fig. 3d) or Student’s *t*-distributions (Fig. 3c, right).

### Maximum likelihood estimation and root-to-tip regression

We generated a maximum likelihood tree using a multiple sequence alignment that included genomes generated in this study, as well as a selection of other available sequences from the Americas, Southeast Asia, and the Pacific. The sequences are listed in Supplementary Information. We ran PhyML^53^ with the GTR substitution model and 4 gamma substitution rate categories; for the tree search operation, we used ‘BEST’ (best of NNI and SPR). In FigTree v1.4.2^54^, we rooted the tree on the oldest sequence used as input (GenBank accession EU545988.1).

We used TempEst v1.5^55^, which selects the best-fitting root with a residual mean squared function, to estimate root-to-tip distances. We performed regression in R with the lm function^47^ of distances on dates. The relationship between root-to-tip divergence and sample dates (Extended Data Fig. 2) supports the use of a molecular clock analysis in this study.

In Supplementary Data, we provide the output of PhyML, as well as the dates and distances used for root-to-tip regression.

### Molecular clock phylogenetics and ancestral state reconstruction

For molecular clock phylogenetics, we made a multiple sequence alignment from the genomes generated in this study combined with a selection of other available sequences from the Americas. We did not use sequences from outside the outbreak in the Americas. Among ZIKV genomes published and publicly available on NCBI GenBank^35^, we selected 32 from the Americas that had at least 7000 unambiguous bases, were not labeled as having been passaged more than once, and had location metadata. We also used 32 genomes from Brazil published in a companion paper^10^ that met the same criteria. The sequences are listed in Supplementary Information.

We used BEAST v1.8.4 to perform molecular clock analyses^56^. We used sampled tip dates to handle inexact dates^57^. Because of sparse data in non-coding regions, we used only the CDS as input. We used the SDR06 substitution model on the CDS, which uses HKY with gamma site heterogeneity and partitions codons into two partitions (positions (1+2) and 3)^58^. To perform model selection, we tested three coalescent tree priors: a constant-size population, an exponential growth population, and a Bayesian Skyline tree prior (10 groups, piecewise-constant model)^59^. For each tree prior, we tested two clock models: a strict clock and an uncorrelated relaxed clock with lognormal distribution (UCLN)^60^. In each case, we set the molecular clock rate to use a continuous time Markov chain rate reference prior^61^. For all six combinations of models, we performed path sampling (PS) and stepping-stone sampling (SS) to estimate marginal likelihood^62,63^. We sampled for 100 path steps with a chain length of 1 million, with power posteriors determined from evenly spaced quantiles of a Beta(alpha=0.3; 1.0) distribution. The Skyline tree prior provided a better fit than the two other (baseline) tree priors (Extended Data Table 2), so we used this tree prior for all further analyses. Using a constant or exponential tree prior, a relaxed clock provides a better model fit, as shown by the log Bayes factor when comparing the two clock models. Using a Skyline tree prior, the log Bayes factor comparing a strict and relaxed clock is smaller than it is using the other tree priors, and it is similar to the variability between estimated log marginal likelihood from PS and SS methods. We chose to use a relaxed clock for further analyses, but we also report key findings using a strict clock.

For the tree and tMRCA estimates in Fig. 2, as well as the clock rate reported in main text, we ran BEAST with 400 million MCMC steps using the SRD06 substitution model, Skyline tree prior, and relaxed clock model. We extracted clock rate and tMRCA estimates, and their distributions, with Tracer v1.6.0 and identified the maximum clade credibility (MCC) tree using TreeAnnotator v1.8.2. The reported credible intervals around estimates are 95% highest posterior density (HPD) intervals. When reporting substitution rate from a relaxed clock model, we give the mean rate (mean of the rates of each branch weighted by the time length of the branch). Additionally, for the tMRCA estimates in Fig. 2c with a strict clock, we ran

BEAST with the same specifications (also with 400M steps) except used a strict clock model. The resulting data are also used in the more comprehensive comparison shown in Extended Data Fig. 3.

For the data with an outgroup in Extended Data Fig. 3, we ran BEAST the same as specified above (with strict and relaxed clock models), except with 100 million steps and with outgroup sequences in the input alignment. The outgroup sequences were the same as those used to make the maximum likelihood tree (see Supplementary Information). For the data excluding sample DOM_2016_MA-WGS16-020-SER in Extended Data Fig. 3, we ran BEAST the same as specified above (with strict and relaxed clocks), except we removed this sample from the input and ran 100 million steps.

We used BEAST v1.8.4 to estimate transition and transversion rates with CDS and non-coding regions. The model was the same as above except that we used the Yang96 substitution model on the CDS, which uses GTR with gamma site heterogeneity and partitions codons into three partitions^64^; for the non-coding regions, we used a GTR substitution model with gamma site heterogeneity and no codon partitioning. There were four partitions in total: one for each codon position and another for the non-coding region (5’ and 3’ UTRs combined). We ran this for 200 million steps. At each sampled step of the MCMC, we calculated substitution rates for each partition using the overall substitution rate, the relative substitution rate of the partition, the relative rates of substitutions in the partition, and base frequencies. In Extended Data Fig. 4, we plot the means of these rates over the steps; the error bars shown are 95% HPD intervals of the rates over the steps.

We used BEAST v1.8.4 to reconstruct ancestral state at the root of the tree using CDS and non-coding regions. The model was the same as above except that, on the CDS, we used the HKY substitution model with gamma site heterogeneity and codons partitioned into three partitions (one per codon position). On the non-coding regions we used the same substitution model without codon partitioning. We ran this for 50 million steps and used TreeAnnotator v1.8.2 to find the state with the MCC tree. We selected the ancestral state corresponding to this state.

In all BEAST runs, we discarded the first 10% of states from each run as burn-in.

In Supplementary Data, we provide BEAST input (XML) and output files. We also provide the sequence of the reconstructed ancestral state.

## Principal component analysis

We carried out principal component analysis using the R package FactoMineR^65^. We imputed missing data with the package missMDA^66^ and we show the results in Fig. 2d.

## Diagnostic assay assessment

We extracted primer and probe sequences from eight published RT-qPCR assays^26–31^ and aligned to our ZIKV genomes using Geneious version 9.1.7^51^. We then tabulated matches and mismatches to the diagnostic sequence for all outbreak genomes, allowing multiple bases to match where the diagnostic primer and/or probe sequence contained nucleotide ambiguity codes (Fig. 3e).

## Data availability

Sequence data that support findings of this study are deposited in NCBI GenBank^35^ under BioProject accession PRJNA344504. Zika virus genomes have accession numbers KY014295-KY014327 and KY785409-KY785485. The dengue virus type 1 genome sequenced in this study has accession number KY829115. See **Supplementary Table 1** for a mapping of sample names to accession numbers.

## Extended Data Figures and Tables

**Extended Data Figure 1.**
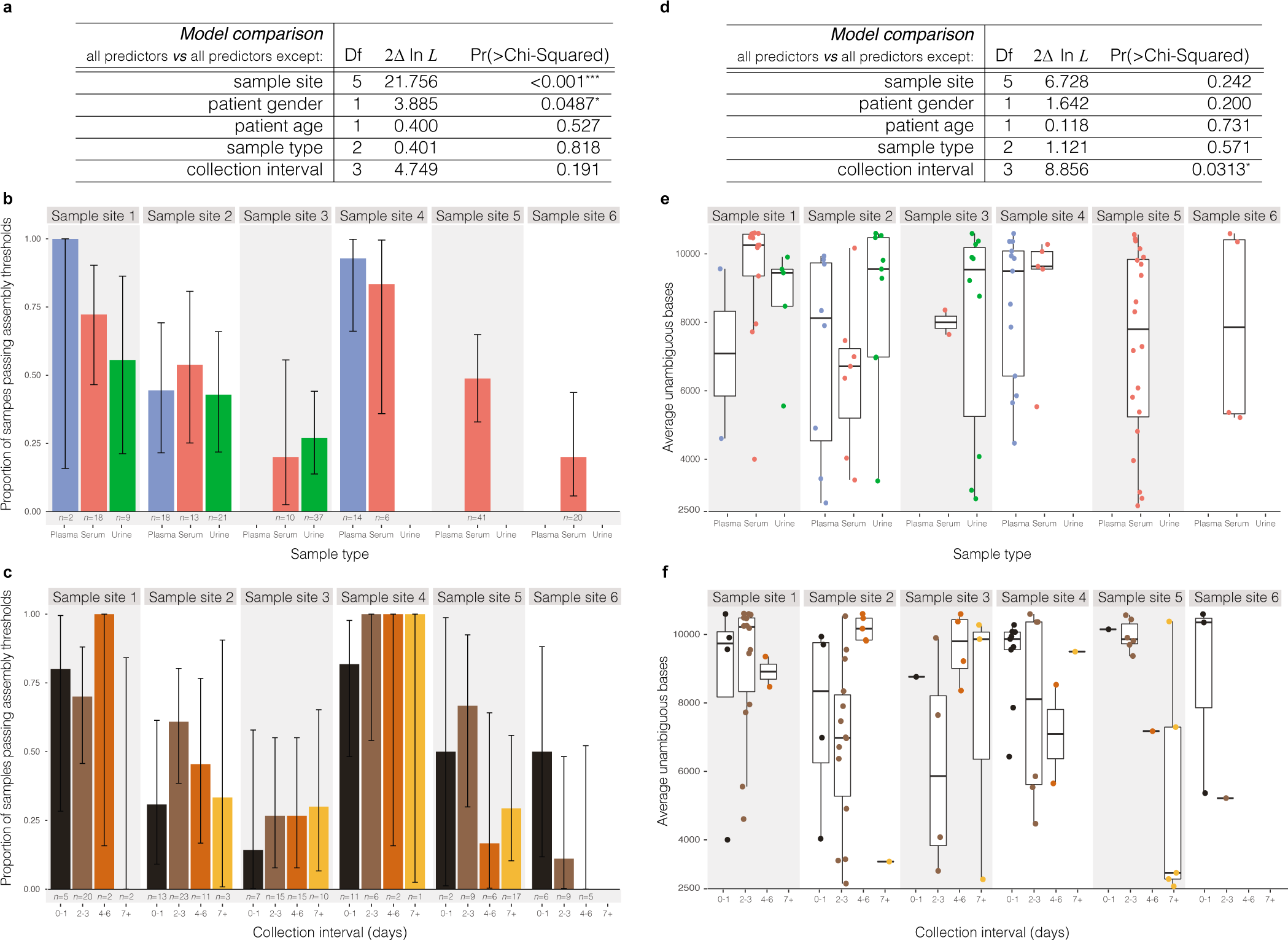
Relationship between metadata and sequencing outcome. Analysis of possible predictors of sequencing outcome: the site where a sample was collected, patient gender, patient age, sample type, and days between symptom onset and sample collection (“collection interval”). **(a)** Prediction of whether a sample passes assembly thresholds by sequencing. Rows show results of likelihood ratio tests on each predictor by omitting the variable from a full model that contains all predictors. Sample site and patient gender improve model fit, but sample type and collection interval do not. **(b)** Proportion of samples that pass assembly thresholds by sequencing, divided by sample type, across six sample sites. **(c)** Same as (b), except divided by collection interval. **(d)** Prediction of the genome fraction identified, using samples passing assembly thresholds. Rows show results of likelihood ratio tests, as in (a). Collection interval improves the model, but sample type does not. **(e)** Sequencing outcome for each sample, divided by sample type, across six sample sites. **(f)** Same as (e), except divided by collection interval. Samples collected 7+ days after symptom onset produced, on average, the fewest unambiguous bases, though these observations are based on a limited number of data points. While the sample site variable accounts for differences in cohort composition, the observed effects of gender and collection interval might be due to confounders in composition that span multiple cohorts. These results illustrate the effect of variables on sequencing outcome for the samples in this study; they are not indicative of ZIKV titer more generally. Other studies^67,68^ have analyzed the impact of sample type and collection interval on ZIKV detection, sometimes with differing results.

**Extended Data Figure 2.**
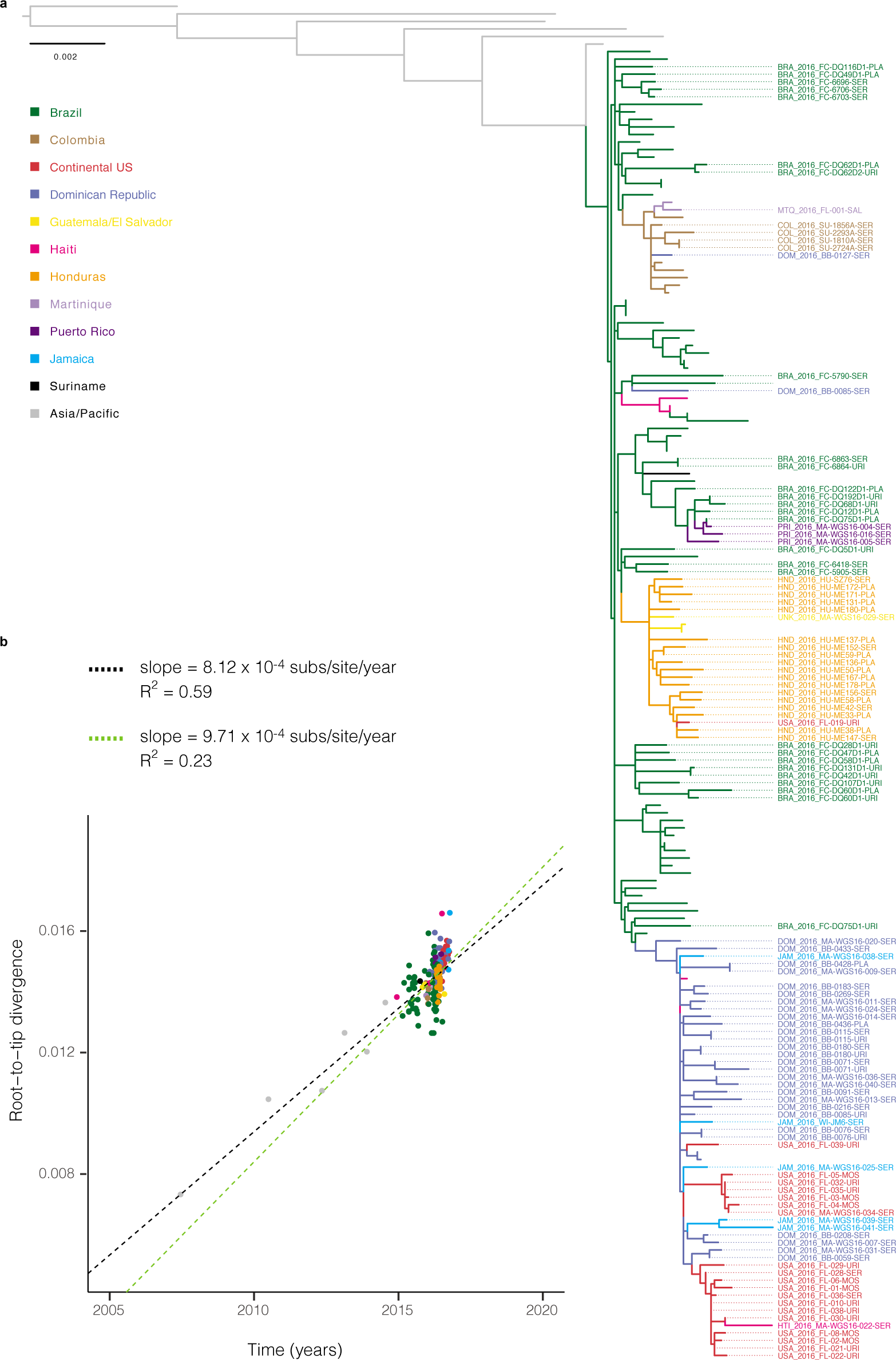
Maximum likelihood tree and root-to-tip regression. **(a)** Tips are colored by sample source location. Labeled tips indicate those generated in this study; all other colored tips are other publicly available genomes from the outbreak in the Americas. Grey tips are samples from ZIKV cases in Southeast Asia and the Pacific. **(b)** Linear regression of root-to-tip divergence on dates. The substitution rate for the full tree, indicated by the slope of the black regression line, is similar to rates of Asian lineage ZIKV estimated by molecular clock analyses^12^. The substitution rate for sequences within the Americas outbreak only, indicated by the slope of the green regression line, is similar to rates estimated by BEAST (1.15×10^−3^; 95% CI [9.78×10^−4^, 1.33×10^−3^]) for this data set.

**Extended Data Figure 3.**
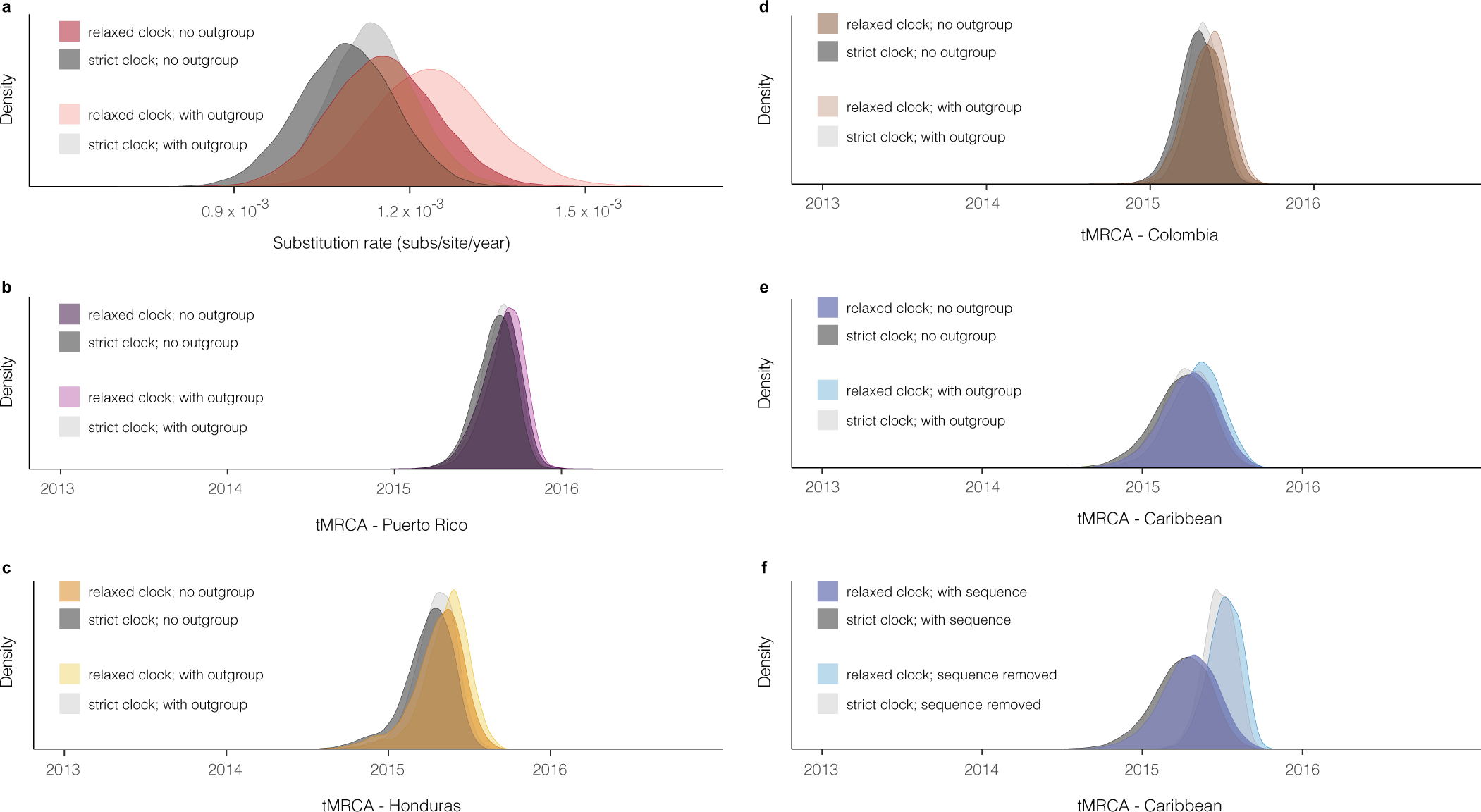
Substitution rate and tMRCA distributions. **(a)** Posterior density of the substitution rate. Shown with and without the use of sequences (outgroup) from outside the Americas. **(b-e)** Posterior density of the date of the most recent common ancestor (MRCA) of sequences in four regions corresponding to those in Fig. 2c. Shown with and without the use of outgroup sequences. The use of outgroup sequences has little effect on estimates of these dates. **(f)** Posterior density of the date of the MRCA of sequences in a clade consisting of samples from the Caribbean and continental US. Shown with and without the sequence of DOM_2016_MA-WGS16-020-SER, a sample from the Dominican Republic that has only 3037 unambiguous bases; this is the most ancestral sequence in the clade and its presence affects the tMRCA. In (a-f), all densities are shown as observed with a relaxed clock model and with a strict clock model.

**Extended Data Figure 4.**
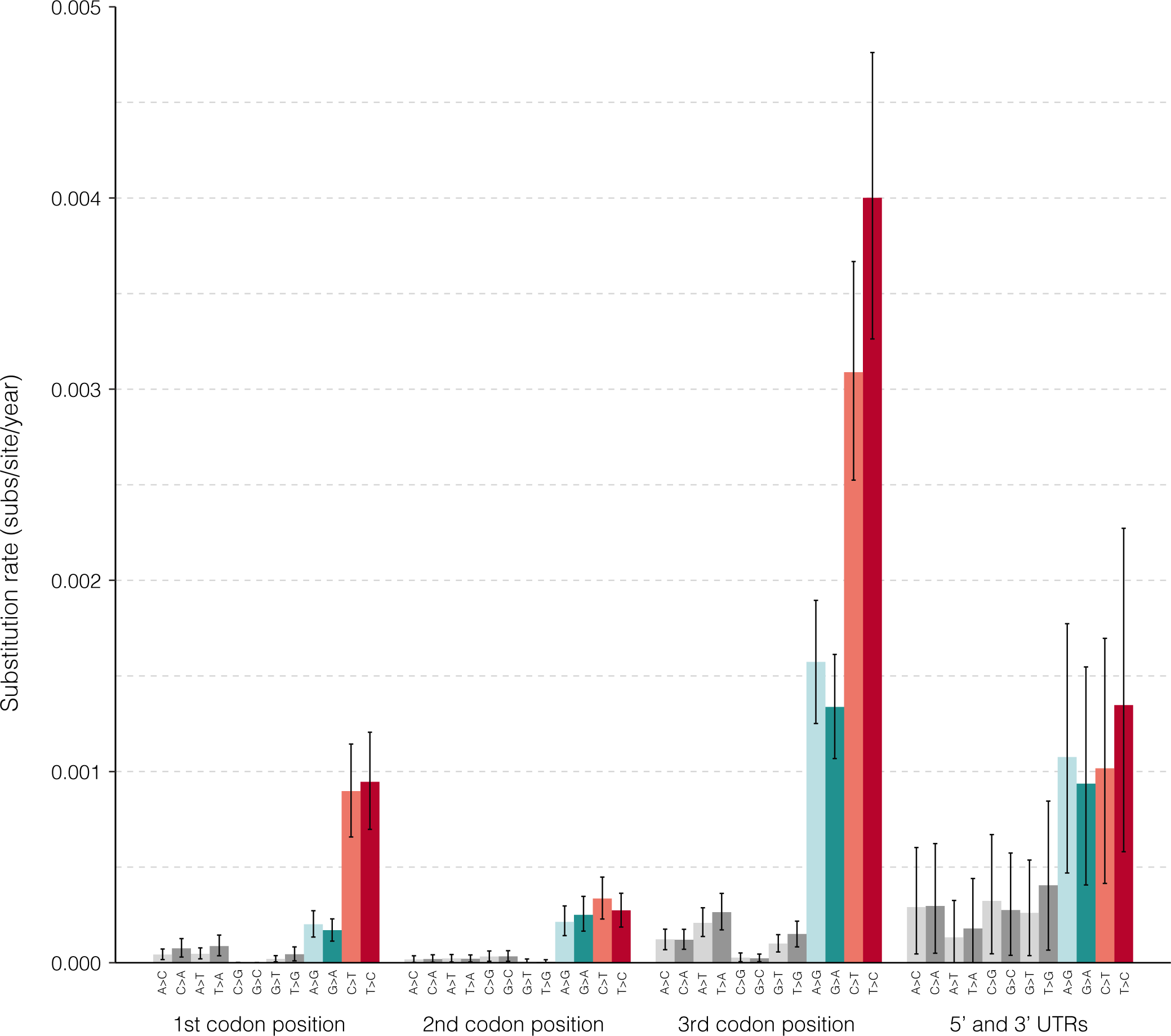
Substitution rates estimated with BEAST. Substitution rates estimated in three codon positions and non-coding regions (5’ and 3’ UTRs). Transversions are shown in grey and transitions are colored by transition type. Plotted values show the mean of rates calculated at each sampled Markov chain Monte Carlo (MCMC) step of a BEAST run. These calculated rates provide additional evidence for the observed high C-to-T and T-to-C transition rates shown in Fig. 3d.

**Extended Data Figure 5.**
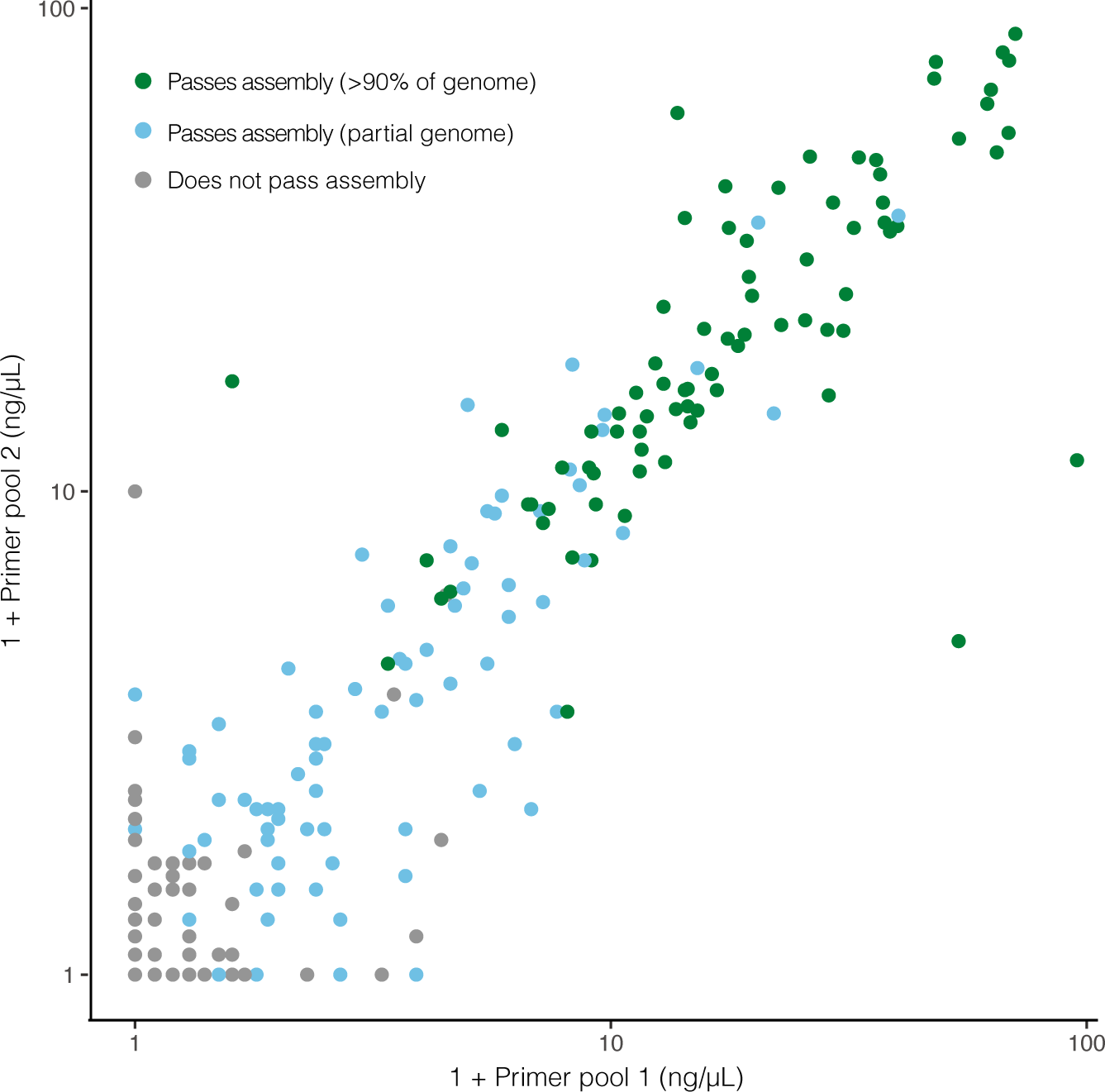
cDNA concentration of amplicon primer pools predicts sequencing outcome. cDNA concentration of amplicon pools (as measured by Agilent 2200 Tapestation) is highly predictive of amplicon sequencing outcome. On each axis, 1+primer pool concentration is plotted on a log scale. Each point is a technical replicate of a sample and colors denote observed sequencing outcome of the replicate. If a replicate is predicted to be passing when at least one primer pool concentration is ≥0.8 ng/μL, then sensitivity=98.71% and specificity=90.34%. An accurate predictor of sequencing success early in the sample processing workflow can save resources.

**Extended Data Figure 6.**
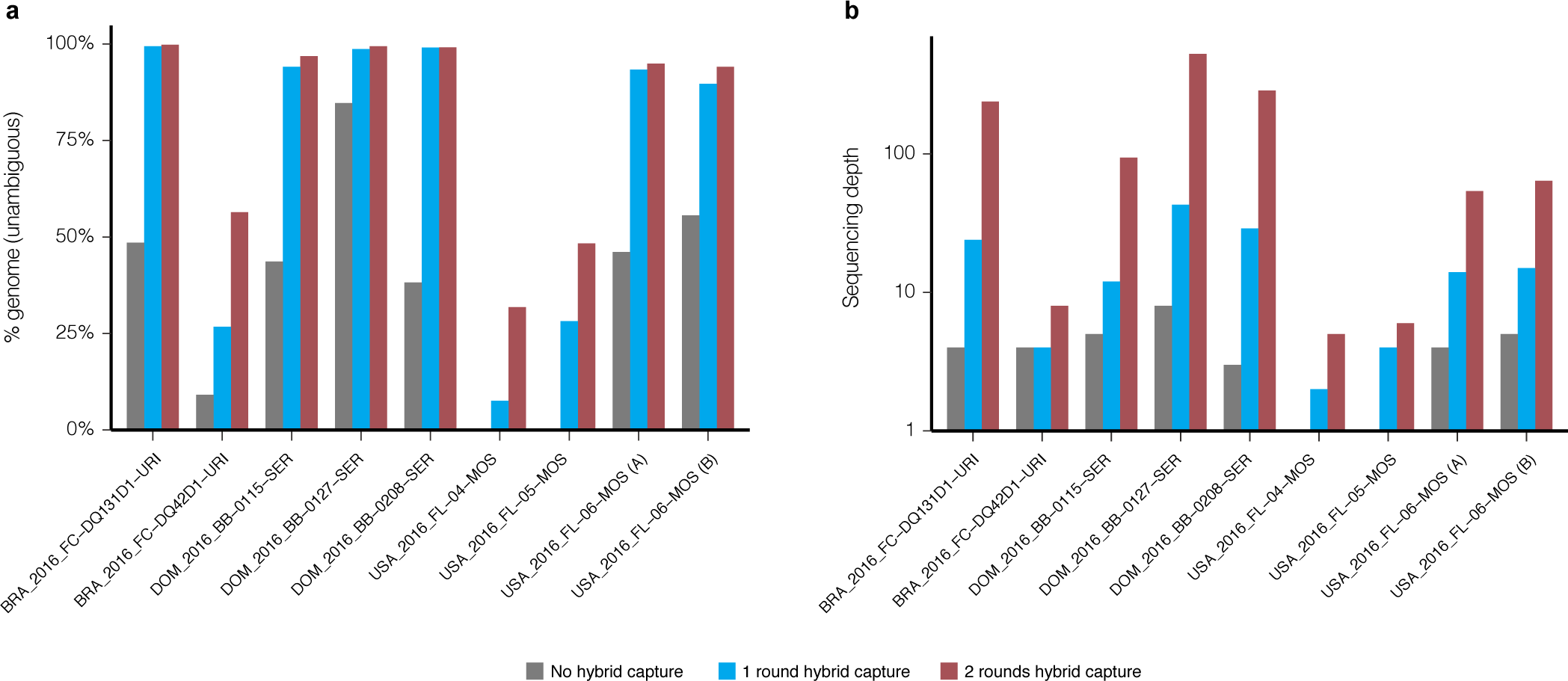
Evaluating multiple rounds of Zika virus hybrid capture. Genome assembly statistics of samples prior to hybrid capture (grey), and after one (blue) or two (red) rounds of hybrid capture. 9 individual libraries (8 unique samples) were sequenced all three ways, had >1 million raw reads in each method, and generated at least one passing assembly. Raw reads from each method were downsampled to the same number of raw reads (8.5 million) before genomes were assembled. **(a)** Percent of the genome identified, as measured by number of unambiguous bases. **(b)** Median sequencing depth of ZIKV genomes, taken over the assembled regions.

**Extended Data Table 1.**
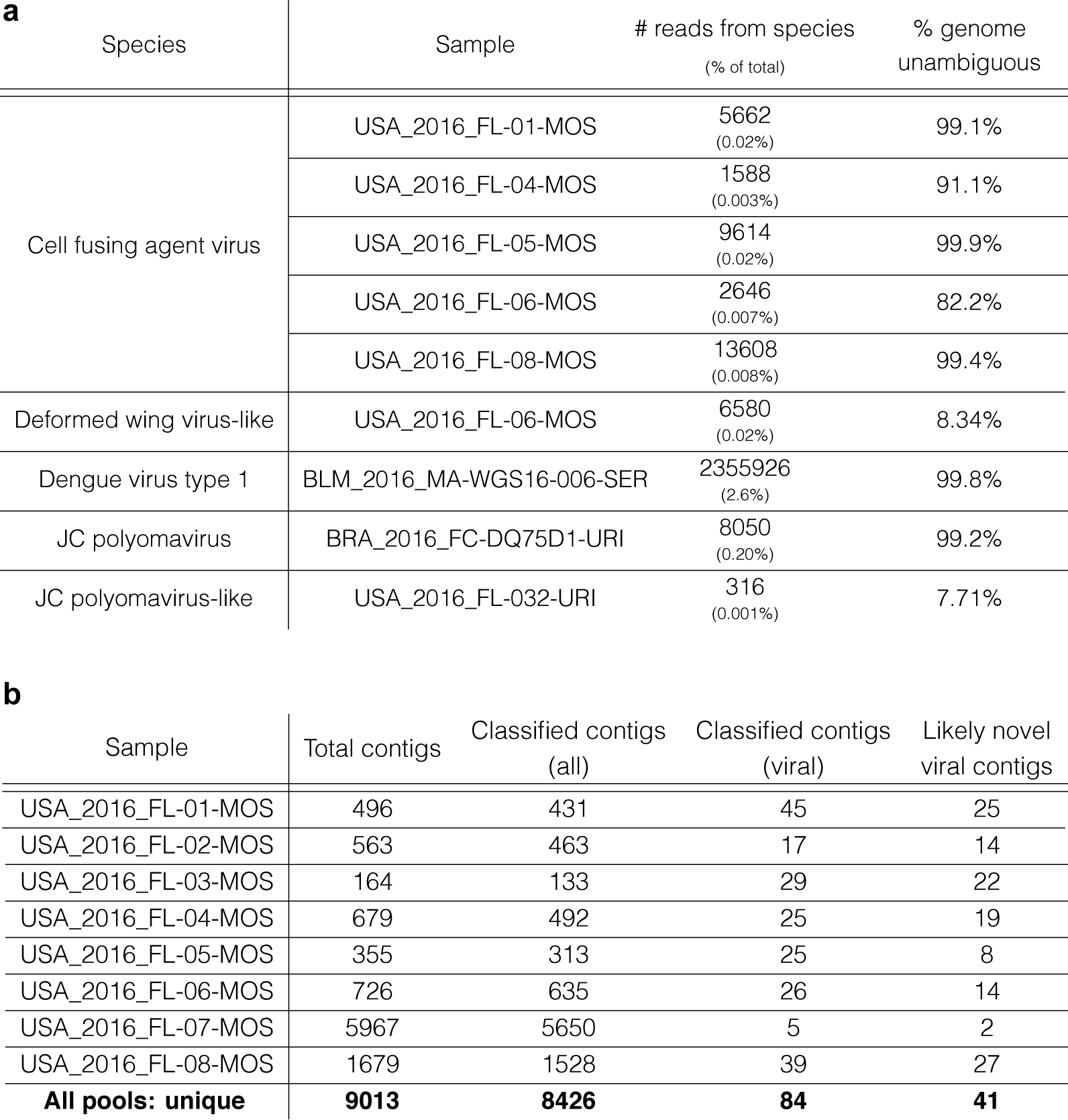
Viruses other than Zika uncovered by unbiased sequencing. **(a)** Viral species other than Zika were found by unbiased sequencing of 38 samples. Column 3: number of reads in a sample belonging to a species as a raw count and a percent of total reads. Column 4: percent genome assembled based on the number of unambiguous bases called. We identified cell fusing agent virus (a flavivirus) and deformed wing virus-like genomes in mosquito pools, and dengue virus type 1, JC polyomavirus, and JC polyomavirus-like genomes in clinical samples. All assemblies had ≥95% sequence identity to a reference sequence for the listed species, except cell fusing agent virus in USA_2016_FL-06-MOS (91%) and dengue virus type 1 in BLM_2016_MA-WGS16-006-SER (92%). The dengue virus type 1 genome showed ≥95% sequence identity to other available isolates of the virus. **(b)** Contigs assembled from unbiased sequencing data of 8 mosquito pools. Column 2: number of contigs assembled. Column 3: number of contigs classified by BLASTN/BLASTX^43^. Column 4: number of contigs hitting a viral species. Column 5: number of contigs hitting a viral species with <80% amino acid identity to the best hit. Each column is a subset of the previous column. Contigs in column 5 are considered to be likely novel. Last row lists counts, after removing duplicate contigs, for all mosquito pools combined. **Supplementary Table 4** lists the unique viral contigs and their best hit.

**Extended Data Table 2.**
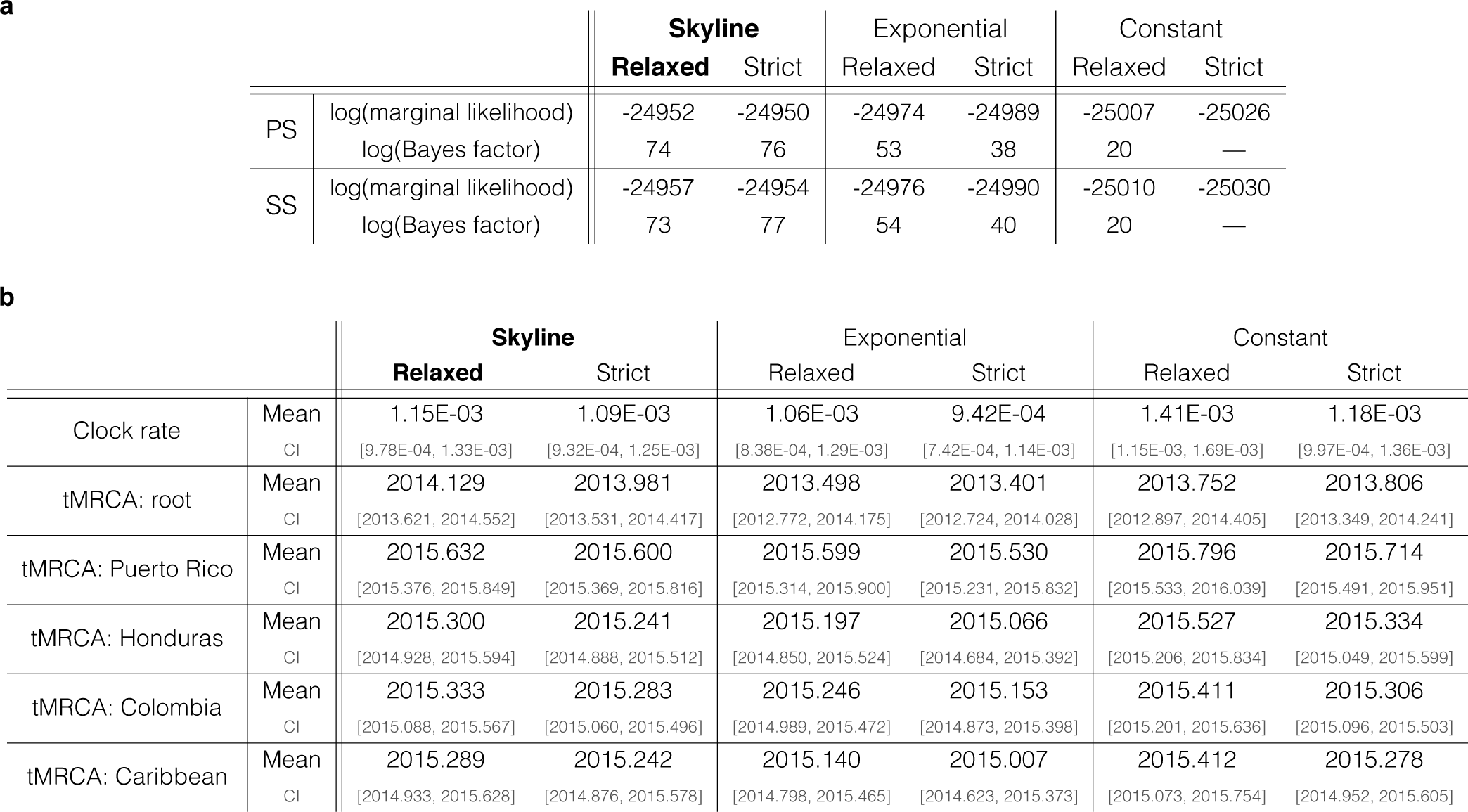
Model selection for BEAST analyses. **(a)** Marginal likelihoods calculated with path sampling (PS) and stepping-stone sampling (SS) for combinations of three coalescent tree priors (constant size population, exponential growth population, and Skyline) and two clock models (strict clock and uncorrelated relaxed clock with log-normal distribution). The Bayes factor is calculated against the baseline model, a constant size tree prior and strict clock. **(b)** Mean estimates and 95% credible intervals (CI) across evaluated models for the clock rate, date of tree root, and tMRCAs of the four regions shown in Fig. 2c. Under a Skyline tree prior, the use of strict and relaxed clock models yields similar estimates.

**a**
|  |  | Skyline |  | Exponential |  | Constant |  |
| --- | --- | --- | --- | --- | --- | --- | --- |
|  |  | Relaxed | Strict | Relaxed | Strict | Relaxed | Strict |
| PS | log(marginal likelihood) | -24952 | -24950 | -24974 | -24989 | -25007 | -25026 |
|  | log(Bayes factor) | 74 | 76 | 53 | 38 | 20 | — |
| SS | log(marginal likelihood) | -24957 | -24954 | -24976 | -24990 | -25010 | -25030 |
|  | log(Bayes factor) | 73 | 77 | 54 | 40 | 20 | — |

**Extended Data Table 2 | Model selection for BEAST analyses.**
|  |  | Skyline |  | Exponential |  | Constant |  |
| --- | --- | --- | --- | --- | --- | --- | --- |
|  |  | Relaxed | Strict | Relaxed | Strict | Relaxed | Strict |
| Clock rate | Mean | 1.15E-03 | 1.09E-03 | 1.06E-03 | 9.42E-04 | 1.41E-03 | 1.18E-03 |
|  | CI | [9.78E-04, 1.33E-03] | [9.32E-04, 1.25E-03] | [8.38E-04, 1.29E-03] | [7.42E-04, 1.14E-03] | [1.15E-03, 1.69E-03] | [9.97E-04, 1.36E-03] |
| tMRCA: root | Mean | 2014.129 | 2013.981 | 2013.498 | 2013.401 | 2013.752 | 2013.806 |
|  | CI | [2013.621, 2014.552] | [2013.531, 2014.417] | [2012.772, 2014.175] | [2012.724, 2014.028] | [2012.897, 2014.405] | [2013.349, 2014.241] |
| tMRCA: Puerto Rico | Mean | 2015.632 | 2015.600 | 2015.599 | 2015.530 | 2015.796 | 2015.714 |
|  | CI | [2015.376, 2015.849] | [2015.369, 2015.816] | [2015.314, 2015.900] | [2015.231, 2015.832] | [2015.533, 2016.039] | [2015.491, 2015.951] |
| tMRCA: Honduras | Mean | 2015.300 | 2015.241 | 2015.197 | 2015.066 | 2015.527 | 2015.334 |
|  | CI | [2014.928, 2015.594] | [2014.888, 2015.512] | [2014.850, 2015.524] | [2014.684, 2015.392] | [2015.206, 2015.834] | [2015.049, 2015.599] |
| tMRCA: Colombia | Mean | 2015.333 | 2015.283 | 2015.246 | 2015.153 | 2015.411 | 2015.306 |
|  | CI | [2015.088, 2015.567] | [2015.060, 2015.496] | [2014.989, 2015.472] | [2014.873, 2015.398] | [2015.201, 2015.636] | [2015.096, 2015.503] |
| tMRCA: Caribbean | Mean | 2015.289 | 2015.242 | 2015.140 | 2015.007 | 2015.412 | 2015.278 |
|  | CI | [2014.933, 2015.628] | [2014.876, 2015.578] | [2014.798, 2015.465] | [2014.623, 2015.373] | [2015.073, 2015.754] | [2014.952, 2015.605] |

**Extended Data Table 3.**
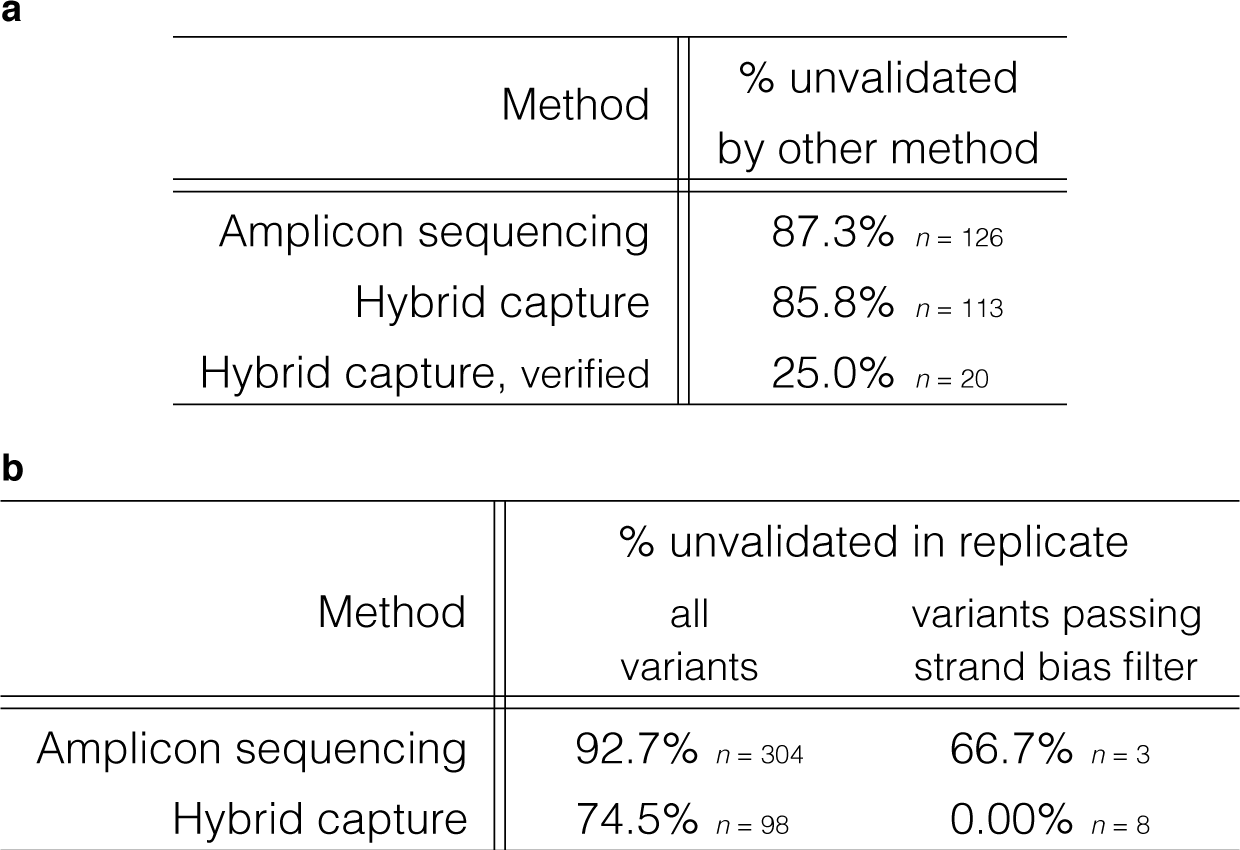
Within-sample variant validation between and within sequencing methods. **(a)** For each method (amplicon sequencing or hybrid capture), fraction of identified variants (≥1%) not identified at ≥1% by the other method (i.e. unvalidated). “Verified” hybrid capture variants are those passing strand bias and frequency filters, as described in Methods. **(b)** For each method, fraction of identified variants unvalidated in a second library. To pass the strand bias filter, a variant must meet filter criteria in both replicates.

**a**
| Method | % unvalidated<br>by other method |
| --- | --- |
| Amplicon sequencing | 87.3% *n* = 126 |
| Hybrid capture | 85.8% *n* = 113 |
| Hybrid capture, verified | 25.0% *n* = 20 |

**Extended Data Table 3 | Within-sample variant validation between and within sequencing methods.**
| Method | % unvalidated in replicate |  |
| --- | --- | --- |
|  | all<br>variants | variants passing<br>strand bias filter |
| Amplicon sequencing | 92.7% *n* = 304 | 66.7% *n* = 3 |
| Hybrid capture | 74.5% *n* = 98 | 0.00% *n* = 8 |

